# A new framework for detecting copy number variants from single nucleotide polymorphism data: ‘rCNV’, a versatile R package for paralogs and CNVs detection

**DOI:** 10.1101/2022.10.14.512217

**Authors:** Piyal Karunarathne, Qiujie Zhou, Klaus Schliep, Pascal Milesi

## Abstract

Studies show that copy number variants (CNVs), due to their ubiquitous presence in eukaryotes, contribute to phenotypic variation, environmental adaptation, and fuel species divergence at a previously unknown rate. However, the detection of CNVs in genomes, especially in non-model organisms is challenging due to the need for costly genomic resources and complex computational infrastructure. Therefore, to provide researchers with a low-cost and easily accessible resource, we developed a robust statistical framework and an R software package to detect CNVs using allelic-read depth from SNPs data.

The core of the framework exploits i) the allelic-read depth ratio distribution in heterozygotes for individual SNPs and testing it against an expected distribution under a binomial sampling, and ii) SNPs showing an apparent excess of heterozygotes under Hardy-Weinberg equilibrium, to detect alleles in putatively multi-copy regions. The use of multiple statistical tests to find the deviation in allelic-read depth ratio distribution makes our method sensitive to sampling and aware of reference biases thereby minimizing false detection of CNVs.

Our framework is well-catered for high throughput short-reads data, hence, most GBS technologies (e.g., RADseq, Exome-capture, WGS). As such, it allows calling CNVs from genomes of varying complexity. The framework is implemented in the R package “rCNV” which effortlessly automates the analysis. We trained our models on simulated data and tested on four datasets obtained from different sequencing technologies (i.e., RADseq: Chinook salmon – *Oncorhynchus tshawytscha*, American lobster – *Homarus americanus*, Exome-capture: Norway Spruce – *Picea abies*, and WGS: Malaria mosquito -*Anopheles gambiae*).

## Background

One main outcome of a decade of massive sequencing efforts is that macro-mutations (e.g., inversions, deletions, insertions, duplications), are prevalent in eukaryotic genomes. Such rearrangements in genomes, known as structural variants (SVs), are crucial for evolution (Chain & Feulner, 2014; Emerson et al., 2008; Fan & Meyer, 2014; Ohno, 1970). When such structural change occurs in an unbalanced process (i.e., insertions, deletions), it leads to copy number variation (CNV) (Collins et al., 2020; Holland et al., 2017). While single nucleotide polymorphism (SNP) is most commonly observed (Savolainen et al., 2013) and used in genetic studies, recent studies found that the number of base pairs affected by structural variants, especially by CNVs is multiple times higher than that by SNPs (Catanach et al., 2019; Sudmant et al., 2013). It means that CNVs can introduce large biases in SNPs calling, and thereby biases in population genetic estimates (Nadukkalam Ravindran et al., 2018; Verdu et al., 2016). On the other hand, CNVs are increasingly acknowledged as a source of genetic polymorphism and for their direct role in an organism’s short-term evolution (Clop et al., 2012; Wellenreuther et al., 2019) and references therein). While most studies considered paralogs (i.e., multiple copies) as difficult to study and hence removed them from analyses, others highlighted CNVs as a source of intraspecific variation during the early divergence of species (e.g. Lauer & Gresham, 2019; Lucek et al., 2019; North et al., 2020; Wellenreuther & Bernatchez, 2018), phenotypic complexity under fluctuating selection: *Mimulus guttatus* – (Nelson et al., 2019), and adaption (Kondrashov, 2012). However, the full spectrum of the role CNVs play in shaping the evolutionary trajectories of species remains grossly unknown (Wellenreuther & Bernatchez, 2018; Wellenreuther et al., 2019).

With the giant leap of high-throughput sequencing technologies (e.g., RADseq, GBS, Exome capture, etc.) millions of genomic sequences allowed researchers to study evolutionary processes at a multi-population scale (Ghosh et al., 2018; Savolainen et al., 2013). However, complex structural variations such as CNVs were overlooked, if not removed, in most population and quantitative genetics analyses (Limborg et al., 2016; McKinney et al., 2017). One main reason for the lack of such studies is the challenge of detecting CNVs in genomes. Several methodologies have been developed to detect CNVs but are mostly restricted to whole genome sequencing (WGS) or whole exome sequencing (WES) data and require well-assembled and annotated reference genomes (Gabrielaite et al., 2021). Such data are expensive to produce impeding large-scale evolutionary studies. This is particularly true for non-model organisms, where reference genomes are often highly fragmented (when available) and poorly annotated. Few recent studies have ventured into this useful type of polymorphism from genotyping by sequencing (GBS) data and demonstrated their advantage when investigating evolutionary changes (e.g., Cayuela et al., 2021; DeBolt, 2010; Dorant et al., 2020; McKinney et al., 2017). For instance, Dorant et al. (2020) called CNVs from RAD-seq data and showed that local adaptation in American lobster (*Homarus americanus*) populations in the southern Gulf of St. Lawrence is better explained by CNVs than by SNP. Similarly, Cayuela et al. (2021) identified CNVs associated with a temperature-dependent clinal shift in the Spotted frog (*Rana luteiventris*) in Columbia. These studies implemented a method originally developed to detect SNPs located in paralogous regions (McKinney et al. 2017) but several crucial aspects of copy number variations inherent to the sequencing technology used were not fully addressed.

Here we present the R software package “rCNV”, a robust framework to identify SNPs located in putatively multi-copy regions; from data pre-processing and filtering to downstream analysis. Our method can be used on any type of short-read data as it considers reference bias and accounts for the errors associated with variation in sample size and depth of coverage. Depending on the downstream analysis, the user can either i) produce a clean dataset free of SNPs lying in putative paralogous or CNV regions for population/quantitative genetic studies and/or ii) call putative CNVs to use them as genetic markers. To do so, we provide further filtering methods combining depth-variation with tests for local enrichment assessments. “rCNV” is a stand-alone comprehensive package that also includes pre-processing functions that allow the handling of raw unfiltered variant call format files (VCF). Downstream analyses using CNVs as genetic markers are also proposed (e.g. *V*_*ST*_, analogous to *F*_*ST*_ (Redon et al., 2006)). The approach implemented in the R package has been optimized to handle a large number of samples and SNPs as well as accounting for sequencing anomalies. The detection methodology is explained in the **Approach** below, and the complete workflow implemented in the ‘*rCNV*’ package is described in the **Analysis** section. The software package is hosted on CRAN [<u>link</u>] and GitHub [<u>link</u>] with a comprehensive tutorial [<u>link</u>].

## Approach

The core of our detection method is a two-step process involving, 1) flagging SNPs deviating significantly from the expected distribution (both mean and variance) of allelic ratio across heterozygous genotypes (i.e., the ratio between the respective read-depth of both alleles in heterozygote individuals) and/or showing an apparent excess of heterozygotes under Hardy-Weinberg equilibrium (HWE) and then 2) identifying SNPs lying in putative multi-copy regions (e.g. CNVs, paralogous, short-tandem repeat) from flagged deviant SNPs.

*Note:* Neither the use of apparent excess of heterozygotes under HWE nor the deviation from expected allelic ratios to detect SNPs as a result of mapping errors or being located in multi-copy regions is new *per se*. For instance, Haldane in 1955 first suggested that “permanent heterozygotes” can be fixed through duplication (see also Spofford 1969, and references therein) and apparent excess of heterozygotes has since been used as a signature for heterogeneous duplication (e.g., Djedatin et al., 2017; Lenormand et al., 1998; Roose & Gottlieb, 1980). Since 2008, allelic ratios have been used to identify the ploidy state of so-called copy number aberrations (Gardina et al., 2008), remove reference bias in short-read data (Gayral et al., 2013; Nabholz et al., 2014), distinguish between sub-genomes in polyploids (Ranwez et al., 2013) or detect SNPs located in paralogous copies (McKinney et al., 2017). As a comprehensive framework, the originality of our approach is to statistically test for the significance of both deviation in expected allelic ratio and higher variance than expected for each SNP independently, taking into account the reference biases (in a broad sense), and variation in the number of heterozygotes and depth of coverage. Finally, our method has been tested and optimized to be used with datasets generated from various short-read sequencing technologies.

### Deviant SNPs (Deviants)

#### 1. Deviation from the expected allelic ratio

In diploid heterozygous genotypes, in the absence of sequencing biases or other error-prone factors, one would expect an equal number of reads supporting each allele and hence a theoretical allelic ratio of 0.5 (see Box 1). This can be seen as the sampling of *N*_*A*_ reads carrying the alternative allele in a binomial distribution of parameter ℬ(*N, p*) where, *N* is the total number of reads covering a given SNP in a heterozygote and, *p*, the probability of sampling a read carrying the alternative allele (*p* = 0.5 in absence of capture bias but can vary between 0 and 1). Therefore, at a given SNP, both the significance of deviation from expected allelic ratios and a higher-than-expected variance in allelic ratios can be tested. Note that the stringency of the tests increases with both read-depths, *N*, and the number of heterozygotes, *n* (see Box 2 and Fig. S1).

#### 2. Excess of heterozygotes under Hardy-Weinberg equilibrium (HWE)

Expected genotype frequencies are computed from observed allele frequencies under HWE (Box. 1). We then perform a Chi-square (*χ*^2^) test to test for the significance of an excess of heterozygote between expected and observed genotype frequencies. We test for an excess of heterozygotes across all samples irrelevant of potential population structure. If population structure were to be present, it should lead to a deficit of heterozygotes (Wahlund effect, Wahlund, 1928). This is rather cautious of false positives.

### Putative multi-copy regions

As mentioned above, factors other than changes in the copy number can explain allelic-ratio deviation (e.g., sequencing technology, mean coverage, quality of the reference genome, methodology used for SNP calling). Hence, not all SNPs flagged as “deviants” may lie in a multicopy region (i.e., because of CNVs or paralogous regions). Therefore, we provide the users with different alternative methods to further filter the dataset to identify putative multi-copy regions;

i. *Intersection set:* In this case, a “deviant” SNP is categorized as located in a putative multi-copy region if it is supported by at least two of the statistics tested above (i.e., excess of heterozygotes, *Z*_*x*_ and χ_x_^2^: see Boxes 1 and 2); see below for our recommendation on using intersection set.
ii. *Unsupervised clustering:* Using *K*-means clustering, the SNPs are grouped into two different clusters (K=2) based on the distribution of *Z*_*x*_, χ_x_^2^, excess of heterozygotes, and read-depth coefficient of variation (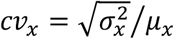 where, 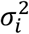 and *μ*_*i*_ are the variance and the mean read-depth of SNP *x* across all individuals).

Note that the *intersection set* approach is based on the significance of the *p*-values for the various scores we computed. The *K-means clustering*, on the other hand, is independent of thresholds or cutoff values as it is conducted on the actual distribution of the statistics and not the *p*-values. For both approaches, users are free to use their own metrics (e.g., allelic probability values) in addition to or in place of the ones already computed. The main purpose of the filtering of putative duplicates from the deviant SNPs is to allow the user to use the best statistics based on their data (see recommendations table in supplementary materials for more details).

#### Box 1

Allelic frequency variation with copy number variation (allele duplication)

**Fig i.**
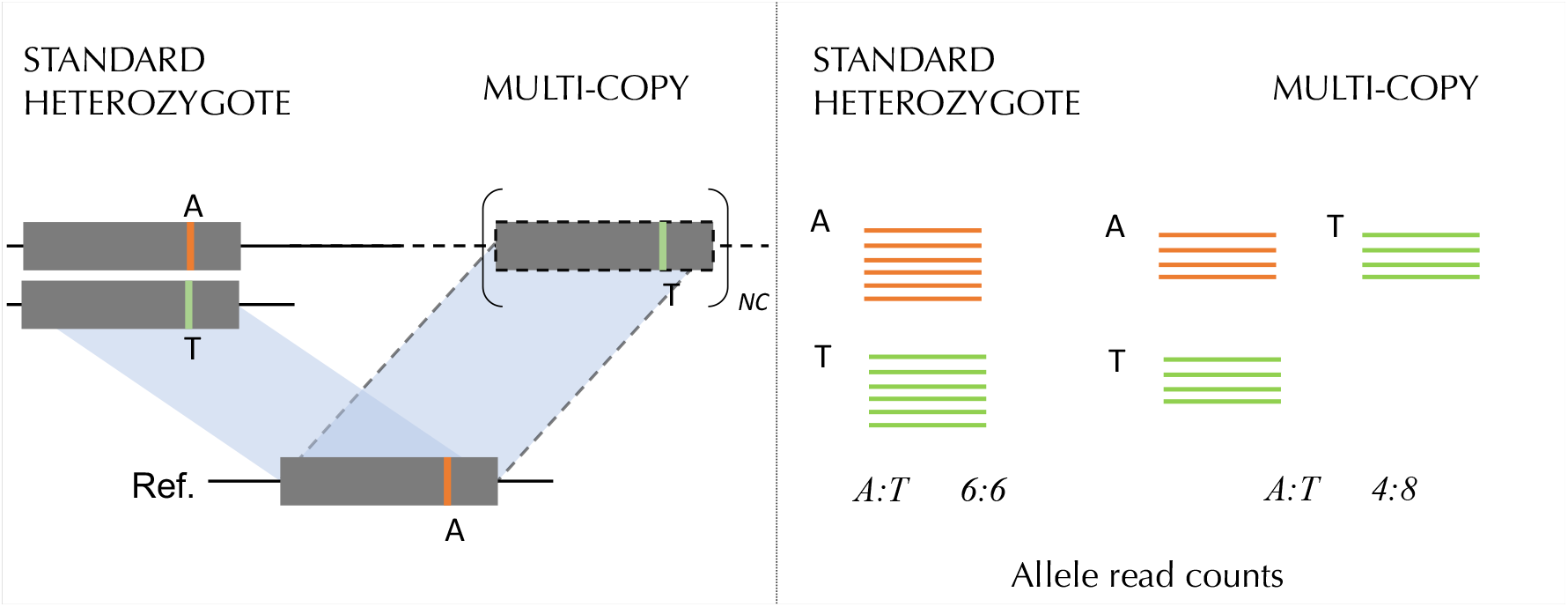
***Schemtic resentation of allele ratio and genotype variation in multi-copy loci compared to a standard heterozygous site in short-read sequencing***.

All the short-reads sequences covering a multicopy region, provided the divergence between the copies is low enough, will be mapped on to the same region of the reference genome, or will be aligned together in reference-free methods, generating “apparent heterozygotes” (i.e., pseudo-SNPs). As a result, one would expect to observe a significant deviation and higher variance in allelic ratios than expected; for “standard heterozygotes,” in absence of external factors (e.g., sequence bias), the expected ratio is 0.5 (Fig.i). It also tends to generate an excess of “apparent heterozygotes” under HWE.

**Fig ii.**
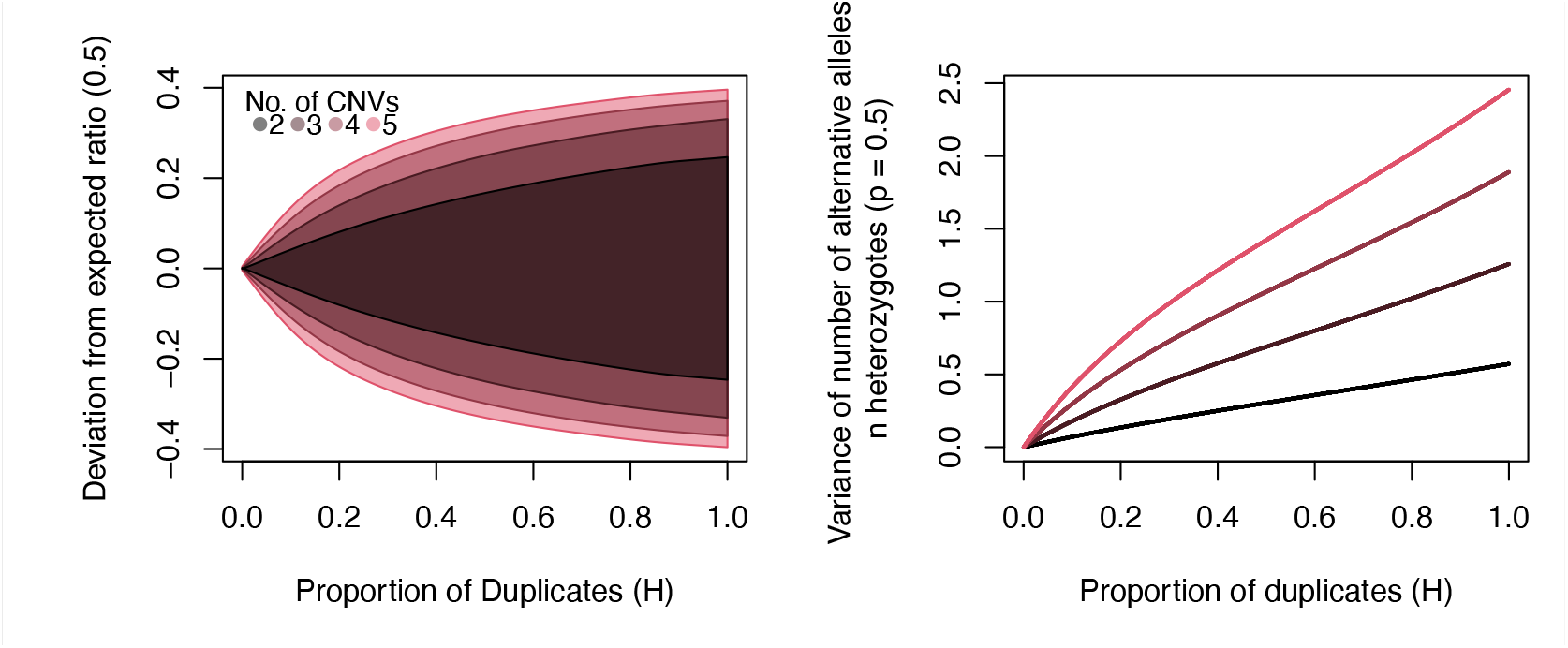
***The deviation of expected allelic frequency in heterozygotes for a) different number of gene copies and proportion of individuals carrying the extra copies and b) the proportion of individuals carrying alternative allele***.

For a given SNP, the distribution of the allelic ratios in a population will therefore depend on, a) the number of copies present, *NC*, b) the proportion of individuals carrying extra copies in the population, *qd* and c) the number of copies carrying the derived allele (Fig.i). Note that all three factors are expected to vary from one SNP to another. Similarly, the number of copies and the proportion of derived allele can vary from one individual to another at a given SNPs. Hence, the deviation from expected allelic ratio in absence of multi-copy region increases with both *NC* and *qd* (Fig.iia). However, when the proportion of derived alleles in the population is close to 0.5, no overall deviation is visible. Nevertheless, the expected variance in number of alternative alleles in heterozygotes also increases with both *NC* and *qd* even for a proportion of derived alleles close to 0.5 (Fig.iib). Equations are presented in Material and Methods “Simulations” and simulations including sampling effect are presented in Figure S1.

#### Box 2

Test for deviation from expected allelic ratio and variance in expected allelic ratio distribution

##### Deviation from expected allelic ratio (Z-score)

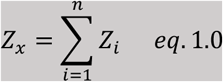

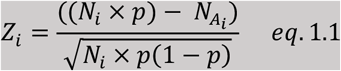

*N*_*i*_ *-total read-depth for heterozygote i at SNP x, N*_*A*_*– alternative allele read-depth for heterozygote i at SNP x, p – probability of sampling allele A for SNP x (p = 0*.*5 if no bias), n – number of heterozygotes at SNP x*.

With *Z*_*i*_ being *n* independent variables following normal distributions, 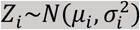, *Z*_*x*_ also follows a normal distribution 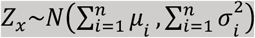. When *N*_A_∼*B*(*N, p*), no deviation from expected allelic ratio is expected and *Z*_*i*_∼*N*(0,1). The significance of the deviation from expected allelic-ratio can thus be tested by computing the quantile *q* = *Z*_*x*_ of the probability density function given by 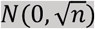.

**Fig ii.**
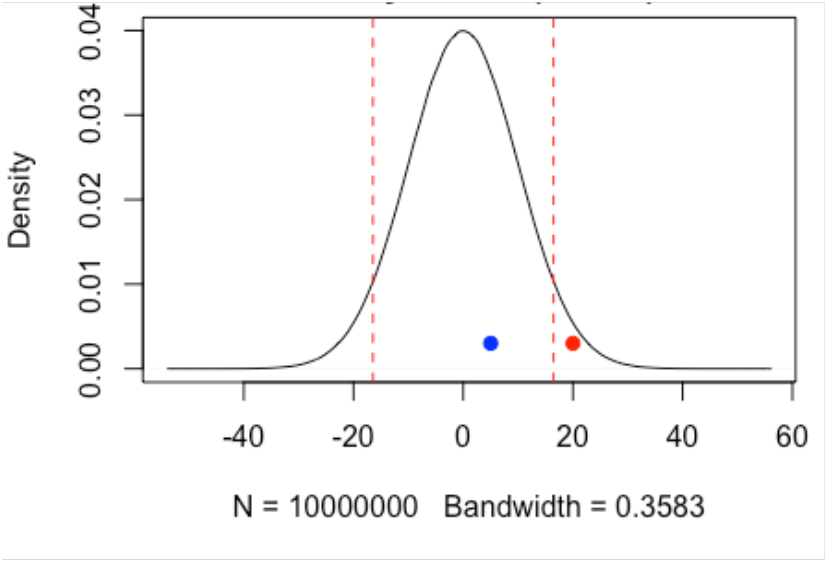
Density distribution of expected *Z*-score. Blue point shows a non-deviant SNP falling inside the significant threshold corresponding to the quantile 0.025 and 0.975 of the probability density function of *N*(0,10). Red point shows a deviant SNP as *Z*_*x*_ correspond to a quantile > 0.975.

##### Deviation from expected allelic ratio (Chi-square)

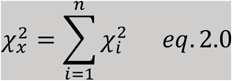

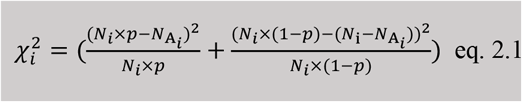

*N*_*i*_ *-total read-depth for heterozygote i at SNP x, N*_*A*_*– alternative allele read-depth for heterozygote i at SNP x, p – probability of sampling allele A for SNP x (p = 0*.*5 if no bias), n – number of heterozygotes at SNP x*.

With 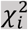 being *n* independent variables following a chi-square distribution with one degree of freedom, 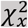 follows a chi-square distribution with *n* degree of freedom. When *N*_*A*_∼*B*(*N, p*), no deviation from expected allelic ratio is expected and the significance of higher variance in allelic-ratio than expected can be tested by computing the quantile 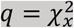 of the probability density function given by 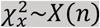.

**Fig.iii.**
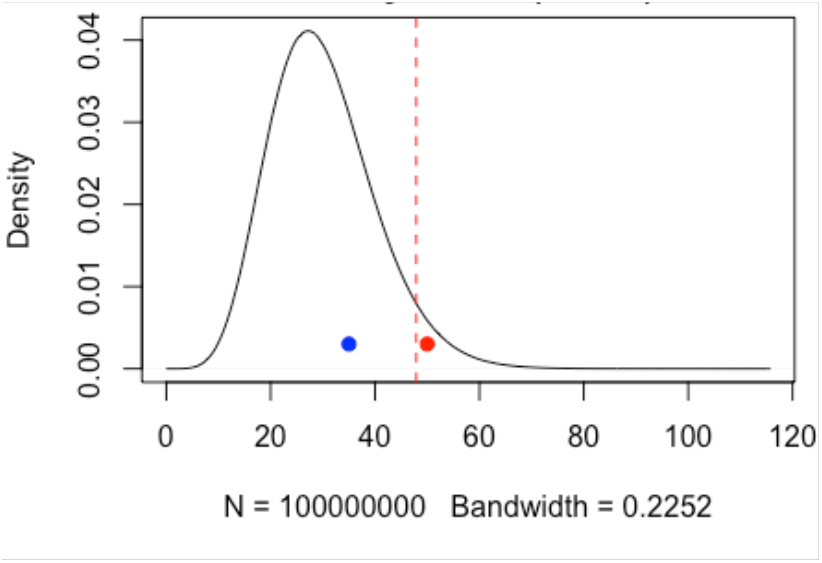
Density distribution of expected *X*^*2*^-score. Blue point shows a non-deviant SNP falling inside the significant threshold corresponding to the quantile 0.025 and 0.975 of the probability density function of *X*(50). Red point shows a deviant SNP as 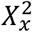 correspond to a quantile > 0.975.

## DATA

In order to evaluate the performance of our approach and showcase the versatility of the *rCNV* package, we used previously published datasets from different sources. The datasets consist of SNPs called from short-reads data acquired through different sequencing technologies (e.g., RADSeq, dd-RADseq, Exome-capture, and WGS) and of varying numbers of samples and SNPs. They are as follows:

### Chinook Salmon

(Larson et al., 2014) – this dataset contains 19,299 SNPs for 266 individuals from five populations. The RADseq-generated data was previously used by McKinney et al. (2017) for paralogs detection. This dataset is particular in the sense that it includes loci located on presumed ongoing residual tetrasomy, hence loci for which the number of copies is known. As in McKinney et al. (2017), we used the annotation made by McKinney et al., (2016) using linkage maps of haploid families to assess the reliability of our detection method.

### American Lobster

(Dorant et al., 2020a) Dryad dataset (Dorant et al., 2020b) – this dataset includes 1,113 lobster samples collected from 21 locations in the southern Gulf of St. Lawrence in Canada, and contains 44,374 SNPs generated using double-digest RAD-seq. After filtering and matching with SNPs used in Dorant et al. (2020) data, we kept 24,193 SNPs from 1,079 samples.

### Norway spruce

(Chen et al., 2019) – *Picea abies* dataset contains over 400,000 SNPs from 453 individuals belonging to 23 populations collected from European natural populations and Swedish breeding programs. The data was generated using exome-capture technology with over 40,000 probes, spread over 20,000 genes. The dataset allowed us to investigate potential biases in allelic-ratio arising from the use of target-capture sequencing technologies.

### Anopheles gambiae

(The-Anopheles-gambiae-1000-Genomes-Consortium, 2017) -*Anopheles gambiae* data set is a subset of 80 samples from the phase 2 AR1 release (https://www.malariagen.net/data/ag1000g-phase-2-ar1) consisting of whole genome sequencing data (Illumina, paired-ends, short read) of 1,142 wild-caught mosquito specimens from 13 countries. Out of the 80 samples we selected, 40 carry an extensive tandem duplication of ∼203Kb (Assogba et al., 2016). Most of the samples carrying the duplication were heterozygotes (i.e., they have a single copy allele on the other strand) and the frequency of the duplication was ∼0.25 considering the whole dataset. We thus used 108,295 SNPs from a 450Kb region centered on the duplication. The use of this dataset is two-fold: i). to show the versatility of our approach using yet another type of sequencing technology and ii). to further test the reliability of the method in detecting CNVs segregating at relatively low frequency.

## ANALYSIS

We followed all the steps of the workflow in Fig. 1 with all the data analyzed with *rCNV* (numbers 1-6 mentioned here in **bold** refer to the steps in the workflow Fig. 1). The raw (unfiltered) VCF files, obtained from the abovementioned databases, were imported to the R environment using the function *readVCF* in rCNV (**1**).

**Figure 1.**
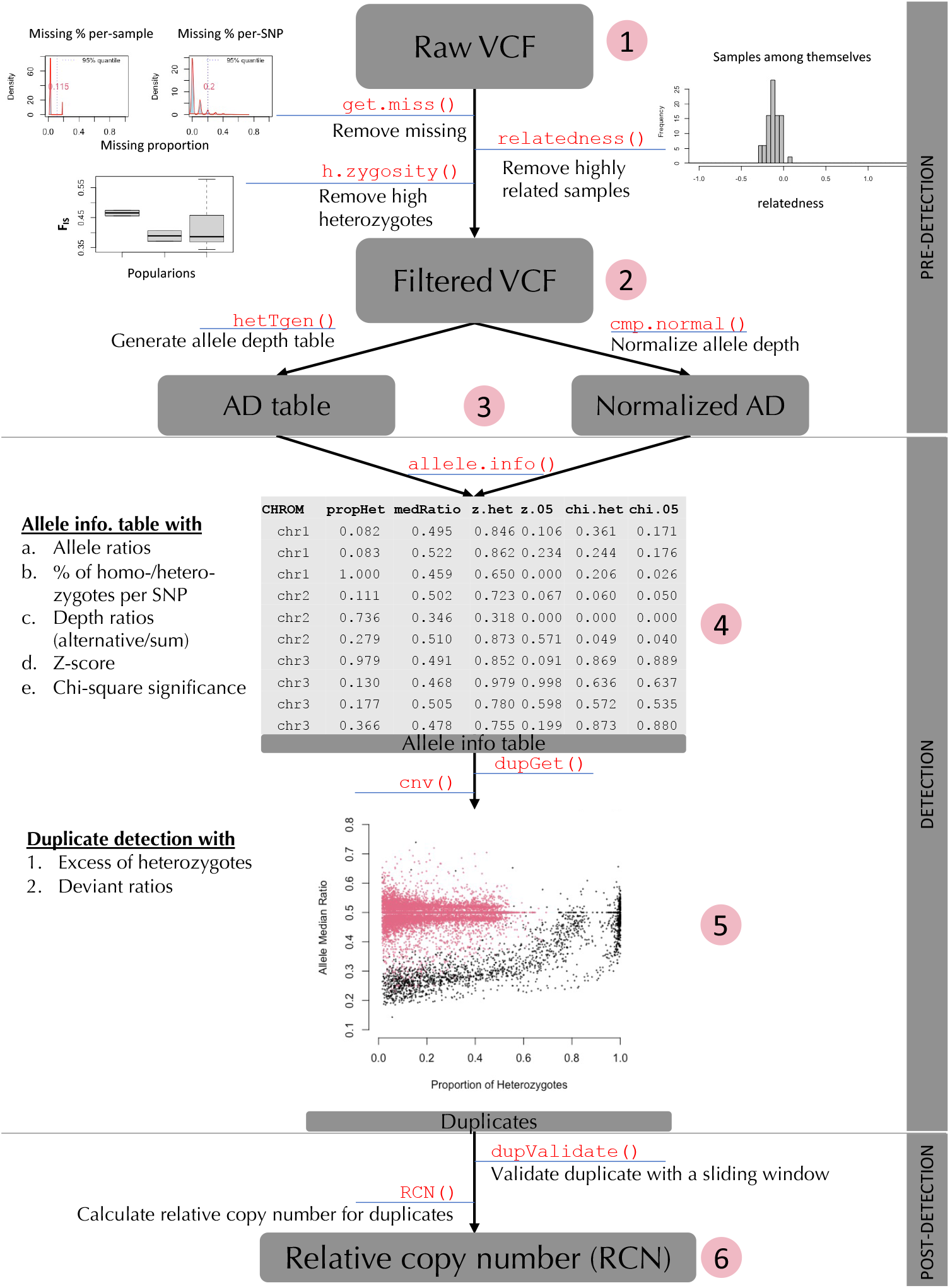
The compact workflow of rCNV, from importing raw data to detecting putative CNV loci and getting the relative copy number, included in the R package.

### 1. Genotypes and variant filtering (not mandatory)

a. Missingness per sample and/or per SNP can be calculated and plotted with *get*.*miss* function to remove low coverage data; user-specific thresholds can be passed on to the function. In our analyses, for all datasets, we kept samples/SNPs for which only less than 30% of data was missing.
b. Individuals’ pairwise relatedness can be calculated according to (Yang et al., 2010) with the *relatedness* function. Individuals with a too-high relatedness can, for instance, represent clones, technical replicates, and siblings but also be sample DNA contamination during library preparation (> 0.9, (Yang et al., 2010). We used a cut-off value of 0.9 for all datasets.
c. Inbreeding coefficient (*F*_*IS*_) can be computed using the function *h*.*zygosity* implementing the following method of moment:

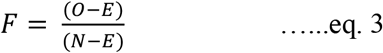

where *O* is the observed number of homozygous sites, *E* is the expected from HWE and N is the total number of sites. Extremely low *FIS* values, often interpreted as a signal for outbreeding, can also indicate potential DNA contamination. We retained all samples above a *FIS* value of -0.2.

*NOTE: If the user has already done VCF filtering to their desired optimization, especially with the above-mentioned parameters, they can start the workflow at number* ***2***.

### 2. Allelic-ratio and metrics computations

a. Filtered VCFs are used to generate tables with depth of coverage values per SNP per allele (**3**). These tables are used as the input to compute various metrics our detection is based on. As the variant calls can contain mismatches between the allele depth (AD) and the genotype (GT) -for instance, the GT called is homozygote (0/0) but AD is non-null for both alleles (e.g. 40:1) -we correct the AD values according to the genotype with *ad*.*correct* function. This function can also be used to correct for the biases introduced by the uneven total depth at a given locus in heterozygote individuals, which introduces additional deviation from the expected allelic ratio. This is done by adding one to the allele with the lowest depth value. Note that by doing so, we are being cautious as we tend to even the allelic ratio.
b. Tables with normalized depth are then generated from the corrected allele depth table (**3**). The aim of the normalization procedure is two-fold: i) checking for potential constitutive bias toward the reference or the alternative allele (i.e., a deviation from expected *p* = 0.5, see box 2) and ii) allowing the quantification of copy number in step **5**. We provide four normalization methods in our package (Median ratio normalization – “MedR”, Quantile normalization – “QN”, Trimmed means of M-values – “TMM”, and PCA-based normalization – “pca”) so that the user can use the optimal normalization method depending on the data and downstream analyses (*cpm*.*normal* function). Normalized depth values are important to investigate the occurrence of probe capture biases or reference mapping bias when assessing allelic ratios, and downstream analyses (e.g., coefficient of variation) as they minimize the biases introduced by the sample effect on the depth of coverage. See the recommendation section for more details on normalization methods. In addition to the normalization of the depth values, *cpm*.*normal* function also flags SNPs and samples with extreme depth values and exports a list of outliers for further filtering.
c. Allele depth table (**3**) is used to calculate several statistics for each SNP independently, among which are, the *Z*_*x*_ and *χ*_*x*_^2^ both of which measure the strength of deviation from the expected allelic ratio (i.e., either mean or variance, box 2), the excess of heterozygotes, and the associated *p*-values (*allele*.*info* function) (**4**). Additional information generated is the proportion of heterozygotes per SNP, the median allelic ratio across all heterozygotes per SNP, and allele frequencies computed either from genotype or depth values.

### 3. Deviant SNPs and putative multi-copy region detection

a. In the last steps of the detection stage (**5**), deviant SNPs are identified based on the significance of the statistics mentioned above (i.e., *Z*_*x*_, *χ*_*x*_^2^ and excess of heterozygotes) with the in-built *dupGet* function. *dupGet* combines deviant ratios and excess of heterozygotes based on the expected allelic ratio; by default, it is 0.5 but it can vary between 0 and 1 and can also be provided by the user. Depending on the technology used for sequencing, some reference bias can exist leading to an over-representation of reads carrying the reference allele, which is often the case for probe-based target capture, for instance. Users are thus recommended to use the expected probe bias as the expected allelic ratio. In addition, depending on the number of SNPs and the distribution of the *p*-values, the user can choose different methods to control for false positive detection propensity associated with multiple testing, see the recommendation section in Table S1 for more details.
b. The SNPs located in putative multi-copy regions can then be identified using one of the two methods presented in the approach section “Putative multi-copy regions” using the *cnv* function. See the recommendation section in Table S1 for more details on using different statistics.
c. If the SNPs are ordered along a reference genome or a genetic map, for instance, the user can adopt a density-based approach using a sliding window to further refine the list of putative SNPs located in putatively multi-copy regions with the *dupValidate* function. Briefly, if the deviation in expected allele ratio and other metrics is caused by the presence of multiple copies at a given locus (*versus* sequencing or technical bias), one would expect several close-by SNPs bearing a signal for the multi-copy region.
d. Finally, a table made of relative copy number is generated from the normalized allele depth table for the putative CNVs and paralogous regions using the *RCN* function. This function is currently under further testing; it uses a PCA-based method to classify the copy number groups based on the total depth values of each locus.

#### Downstream analyses

We have implemented the comparative statistic *V*_*ST*_ (variance fixation index – Redon et al., 2006), calculated from depth values of putative CNV loci to assess the genetic variation among populations. *V*_*ST*_ is analogous to fixation index (*F*_*ST*_) in population genetic analyses, and calculated as *V*_*ST*_ = 1 − *V*_*S*_/*V*_*T*_ where *V*_*T*_ is the variance of normalized read depths among all individuals from two populations and *V*_*S*_ is the average of the variance within each population, weighed for population size (Dennis et al., 2017; Weir & Cockerham, 1984). We further provide the option to plot a network graph to visualize the among-sample association based on the *V*_*ST*_ distance matrix. The high-confident putative CNVs filtered through post-detection are then used to calculate the relative copy number (RCN) across samples per locus (see prospects) and can be used in further downstream analyses such as genome-wide association analysis (GWAS) and environmental association analysis (EAA).

## SIMULATIONS

The use of simulation is twofold: 1). investigating the patterns of deviation from expected allelic-ratio due to i) sampling size, ii) number of copies at a given locus and the proportion of derived alleles, and iii) frequency of the duplicated alleles in the dataset, and 2) assessing the power of the statistical test using *Z*_*x*_ and 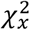 statistics.

### 1- Patterns of deviation in allelic ratio

#### Sampling size

The expected number of reads covering each allele in a heterozygous genotype should follow a binomial distribution of parameter ℬ(*N, p*) where *N* is the total number of reads covering a given SNP, and *p* is the probability of sampling a read carrying a given allele (*p* = 0.5 in absence of bias). When several heterozygous genotypes are found at the same position, this sampling is repeated *n* times, *n* being the number of heterozygote individuals for that position, ℬ(∑_*n*_ *N, p*) . Besides sequencing error, reference bias, or the presence of multi-copy regions, some deviation from *p* can be expected simply because of the sampling effect, especially when *N* and/or *n* is small. To illustrate this, we performed simulations by randomly sampling *n* times in a binomial distribution ℬ(*N*, 0.5) using the *rbinom* function of the *stats* R package (v4.1.0, Kachitvichyanukul & Schmeiser, 1988). We then computed the absolute average deviation from 0.5 for each combination of *n* and *N* parameters (both ranging from one to 100 with an increment of one) and averaged it across 10,000 simulations (*sim*).

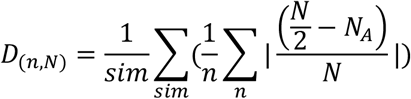

With *N*_*A*_, a vector of length *n* of random sampling in a binomial distribution ℬ(*N*, 0.5).

#### Number of copies and frequency

We further illustrated the deviation in allelic ratio induced by the variable number of copies in the duplicated allele and their frequencies in the population. For a given number of copies in a duplicated allele, *NC*, the number of copies of individuals carrying *i* duplicated allele (*i* can be 0, 1, or 2 for diploids), *m*_*i,NC*_, is given by:

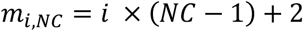

The number of individuals carrying *m*_i,NC_ copies is thus:

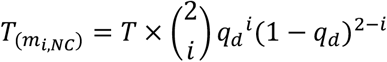

Where *q*_d_ is the frequency of the duplicated allele in the population and *T* is the total number of individuals in the population. To simplify the simulation, we assumed that the alternative allele frequency, *p*, is uniform across duplicated and non-duplicated alleles; *i*.*e*., the number of copies carrying the alternative mutation is proportional to the frequency of the alternative mutation in the whole population. Based on Hardy-Weinberg equilibrium, the number of heterozygous genotypes with *k* alternative copies for a given *m, T*_(k,m)_, can be derived from a binominal sampling:

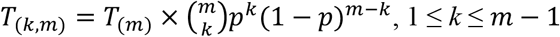

The expected allelic ratio for each heterozygous genotype for a given *m, R*_(*k,m*)_, can be calculated as:

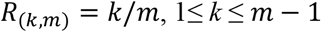

The expected average allelic ratio deviation in the population, *E*_*D*_, given that the expected allelic ratio in a single copy region of 0.5, is therefore:

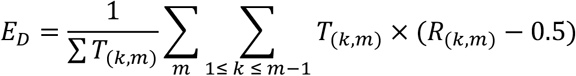

It is worth noting that *E*_*D*_ will fail to grasp deviation when the distribution of *p* is centered around 0.5. However, the expected variance in the number of alternative alleles in heterozygotes, *E*_*V*_, increases along with both *NC* and *q*_*d*_, and is given by:

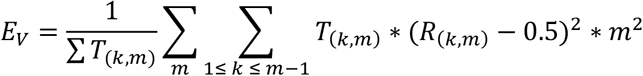

We then simulated a population with 100 individuals. To incorporate the sampling effect, the sequencing coverage of individual *j* with *m* copies was randomly sampled in a normal distribution *N*_(*j*)_∼*N*(20 ∗ *m*, 10). The number of copies of a duplicated allele, *NC*, ranged from one to five. The frequency of a given duplicated allele, *q*_*d*_, and of the alternative mutation, *p*, in the population varied from zero to one in 0.01 increments. For each combination of *NC, q*_*d*_, and *p*, the average allelic ratio deviation, *D*, and variance of the number of alternative alleles, *V*, were calculated as:

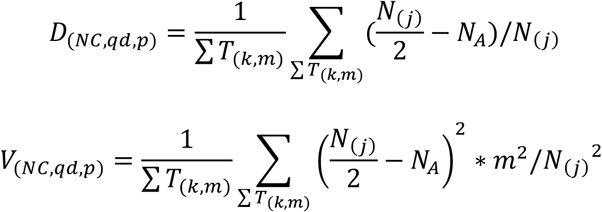

Where *N*_*A*_ is a random draw from ℬ(*N*_(*j*)_, *R*_(*k,m*)_) given the number of copies, *m*, and the number of alternative mutations, *k*, carried by the heterozygote *j. p* was fixed to 0.5 when calculating *V*.

### 2- Power analyses

We assessed the power of the detection test using the *Z*_*x*_ and 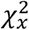 statistics depending on the depth, *N*, and the number of heterozygotes *n*, as well as a function of the intensity of the deviation in allelic ratio, *p*. We used the same simulation framework as above for illustrating the sampling effect. To simulate a consistent bias toward a given allele, *p* ranged from 0 to 0.5 in 0.005 increments. For *Z*_*x*_, *N*_*A*_ is a vector of length *n* of random sampling in a binomial distribution ℬ(*N, p*). For 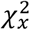, *N*_*A*_ is a vector of length *n* with *n/2* random sampling in binomial distribution ℬ(*N, p*) and ℬ(*N*, (1 − *p*)). For each combination of *n, N* (both ranging from one to 100 by steps of one), and *p* parameters, we computed the *Z*_*x*_ and 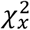 statistics as presented in Box. 2 considering an expected allelic ratio of 0.5. The significance of the deviation from the expected allelic ratio was then computed as the quantile of q = *Z*_(*N,n,p*)_ of the probability density function given by 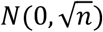 and the significance of higher variance in allelic-ratio than expected can be tested by computing the quantile 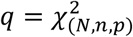 of the probability density function given by 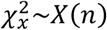.

The whole procedure was repeated 10,000 times for each *N, n*, and *p* parameter combinations and independently for *Z*_*x*_ and 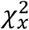. We then recorded the number of significant deviations (*q* < 0.05). Figures S1B and C show the 95% true positive detection rate for the various depth, number of heterozygotes, and intensity of deviation from expected allelic – ratio respectively for *Z*_*x*_ and 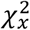. The user can test this using the function *power*.*bias()* in the package.

## Results

All the VCF files for each of the four datasets were filtered (initial variant filtering) for SNPs with a low number of heterozygotes (< 4 heterozygotes per SNP) and outlier samples and SNPs where extreme depth values were recorded. All the removed SNPs had less than 10% heterozygotes in all the datasets. Such low levels of heterozygosity are uninformative and can negatively affect the CNVs detection (see Approach), and therefore were removed from our analyses. The SNPs with a median read depth of < 5 were also not considered for CNV classification and were removed from all the analyses (classified as low confidence in simulations: Fig. S1A).

### Chinook salmon

Out of 19,299 SNPs, only 18,309 were kept for analysis after initial variant filtering. Out of these SNPs, 14.8% were identified as *deviant* SNPs, of which 11.3% were recognized as located in putative multi-copy regions. Because of the recent tetrasomy, an allelic ratio of up to 1:3 is expected for SNPs located in the tetratomic regions. This is what we observed, with a clear definition in the allele ratio plot (Fig. 2) between the SNPs located in single-copy regions (pink) and those that are in multi-copy regions (green & blue). Out of all the common SNPs (8,220) analyzed in both our analysis and McKinney et al. (2016), the latter identified 1,223 (14.87 %) SNPs as so-called “duplicates” while *rCNV* detected 1,171 (14.24 %) as putatively located in multi-copy regions (Table 1 & Fig.S2A). Assuming that the detection of paralog in McKinney et al. 2016 is the true paralogs, our detection using *rCNV* is 96% accurate. From 6,997 so-called “singletons” detected by McKinney et al. 2016, *rCNV* detected 6,983 as non-deviant SNPs, showing 99.15% accuracy for non-CNV classification. We labeled SNPs as pseudo false positives (*pFP*) or pseudo false negatives (*pFN*) if the classification by *rCNV* does not match that of McKinney et al. 2016 (Figs S2B&C). The *pFP*, classified as putatively located in multi-copy regions by *rCNV* but classified as non-duplicated in McKinney et al. 2016, have allelic ratios expected for the tetratomic region (i.e., 3:1 or 1:3). On the other hand, *pFN* (i.e., classified as duplicated in McKinney et al. 2016 analysis but classified as non-deviant by *rCNV*) are in the low confidence zone (i.e., where either the number of heterozygotes is low e.g., 10 or less per SNP, or the median depth values are on the low side for confident detection (e.g., <6), as shown in simulations Fig. S1).

**Figure 2.**
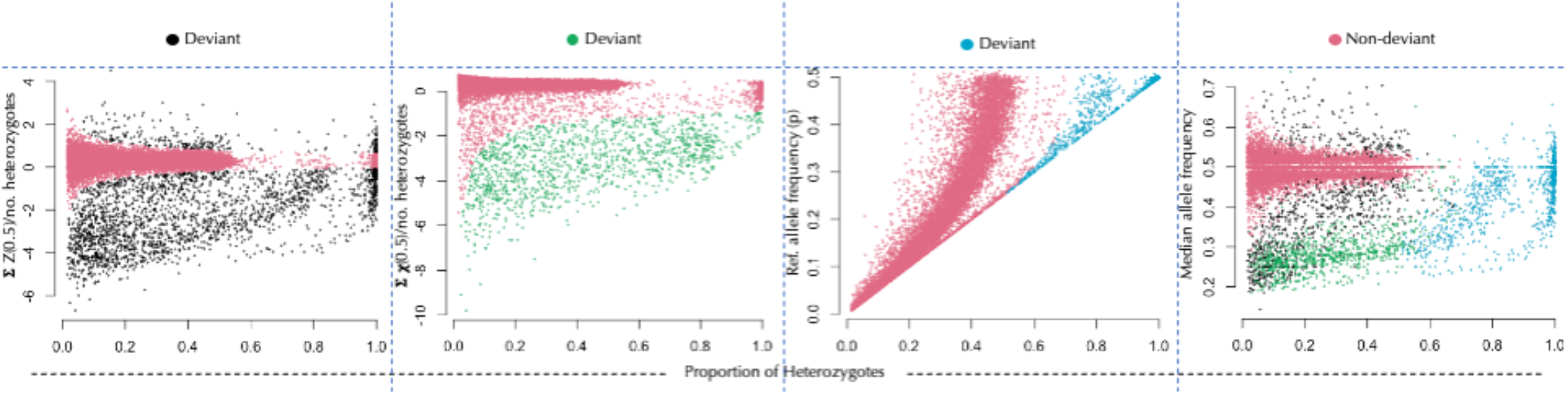
Different ranges of deviant SNP detection by (plots left to right) Z-score distribution of depth values, chi-square distribution of depth values, excess of heterozygotes, and the combination of all statistical methods on Chinook Salmon data (Larson et al., 2014).

**Table 1.** Copy number variation detection by rCNV for different datasets with their respective sequencing technology, size (i.e., samples & SNPs), deviant SNPs, putative CNVs, and overlap with previous studies.

| Species | Technology | # |  | Deviants |  | CNVs |  | Overlap†<br>(%) |
| --- | --- | --- | --- | --- | --- | --- | --- | --- |
|  |  | Samples | # SNP | # | % | # | % |  |
| Chinook Salmon | RAD-Seq | 226 | 18,309 | 2,717 | 14.8 | 2,070 | 11.3 | 96 |
| American Lobster | ddRAD-Seq | 1,074 | 24,047 | 7,125 | 29.6 | 3,596 | 14.95 | 41 |
| Norway Spruce | Exome capture | 423 | 245,758 | 16,064 | 6.5 | 5,397 | 2.19 | - |
| *Anopheles gambiae* | WGS | 80 | 16,471 | 759 | 4.6 | 565 | 3.4 | - |

#: number; †: overlap compared to respective previous study

### American lobster

After filtering, *rCNV* only kept 24,047 SNPs for duplicate detection. In the American lobster dataset, we flagged 7,125 (29.6%) SNPs as *deviants*, of which we detected 3,596 SNPs (14.95%) as putatively located in multi-copy regions, whereas Dorant et al. (2020) identified 39.92% (9,659 out of 24,193) to be located in duplicated regions (Table 1 and Fig. S3). Out of all SNPs detected as putatively duplicated by either analysis (8,162 SNPs), 41% were detected in both analyses, but only 64 SNPs were detected as located in putatively CNV regions with *rCNV* that were not detected in Dorant et al. analysis, here again, we demonstrate a higher stringency in our detection approach; whereas their study detected additional 4,593 alleles that we did not detect. We labeled these as pseudo false positives (*pFP*: detected as putative CNVs by rCNV but classified as non-duplicated by Dorant et al.) and pseudo false negatives (*pFN*: classified as duplicated by Dorant et al. and non-CNVs by rCNV): *pFP* – 64, pFN – 4,593. To visualize this overlap (and the lack thereof), we plotted *pFP* and *pFN* on allele ratio plots (Fig. S4). Almost all the *pFN* detection (purple points in the plot) by *rCNV* are either in the low-heterozygosity zone (mean 15 individuals) or have an allelic ratio close to 1:1 (blue cloud), and they were not detected as deviants in our analysis with any of the three statistics tested, likely indicating high false positive detection rates in Dorant et al. (2020).

### Norway spruce

After initial filtering, 245,758 SNPs were kept for analysis. Out of which, 10.51 % (25,826 SNPs) were flagged as *deviants*. Among them, 4,201 SNPs (1.71%) were located in putatively CNV regions (Fig. S5). Comparing mean read coverage between homozygotes for the reference or the alternative allele indicated the presence of a large sequencing probe bias towards the reference allele (Fig. S6). Therefore, at a given SNP, we used the average allelic ratio (i.e., the ratio between the average coverage of the alternative allele and the sum of the average coverage of both alternative and reference alleles, using normalized depth) computed across all samples as the expected allelic ratio (instead of *p* = 0.5 in eq. 1.1 and 2.1 in Box. 2) for computing both *Z*_*x*_ and 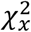 statistics (*z*.*all* and *chi*.*all* in the functions *dupGet* and *cnv*). We tested the gain in accuracy by comparing the distribution of depth distribution and local clustering with that of the classification using *z*.*05* and *chi*.*05*, thereby assuming no probe sequencing bias.

### Anopheles gambiae

The target region (400 kb) of chromosome arm 2 (2R) with the known duplicated region consisted of ∼145,000 SNPs after filtering out multi-allelic loci. Out of this, only 16,471 SNPs were kept for the analysis after filtering for low number of heterozygotes and low depth of coverage. We found that 565 SNPs bear a signal for being located in the putatively duplicated region (Fig. S7) among the 759 that were classified as deviant SNPs. While most of the SNPs (68%) detected as putative CNVs are actually in the region *AgamP4_2R:3436927-3639836*, which was previously described as a duplicated chromosome fragment (Assogba et al., 2016), we also found SNPs that are putative CNVs in the flanking the regions of *AgamP4_2R:3436927-3639836* (Fig. 3A); this observation is not surprising as the flaking regions are rich in transposable elements (Assogba et al., 2016). However, out of all the SNPs in the putatively duplicated region (excluding the SNPs in the flanking regions), 84% were detected as putative CNVs in our analysis.

**Figure 3.**
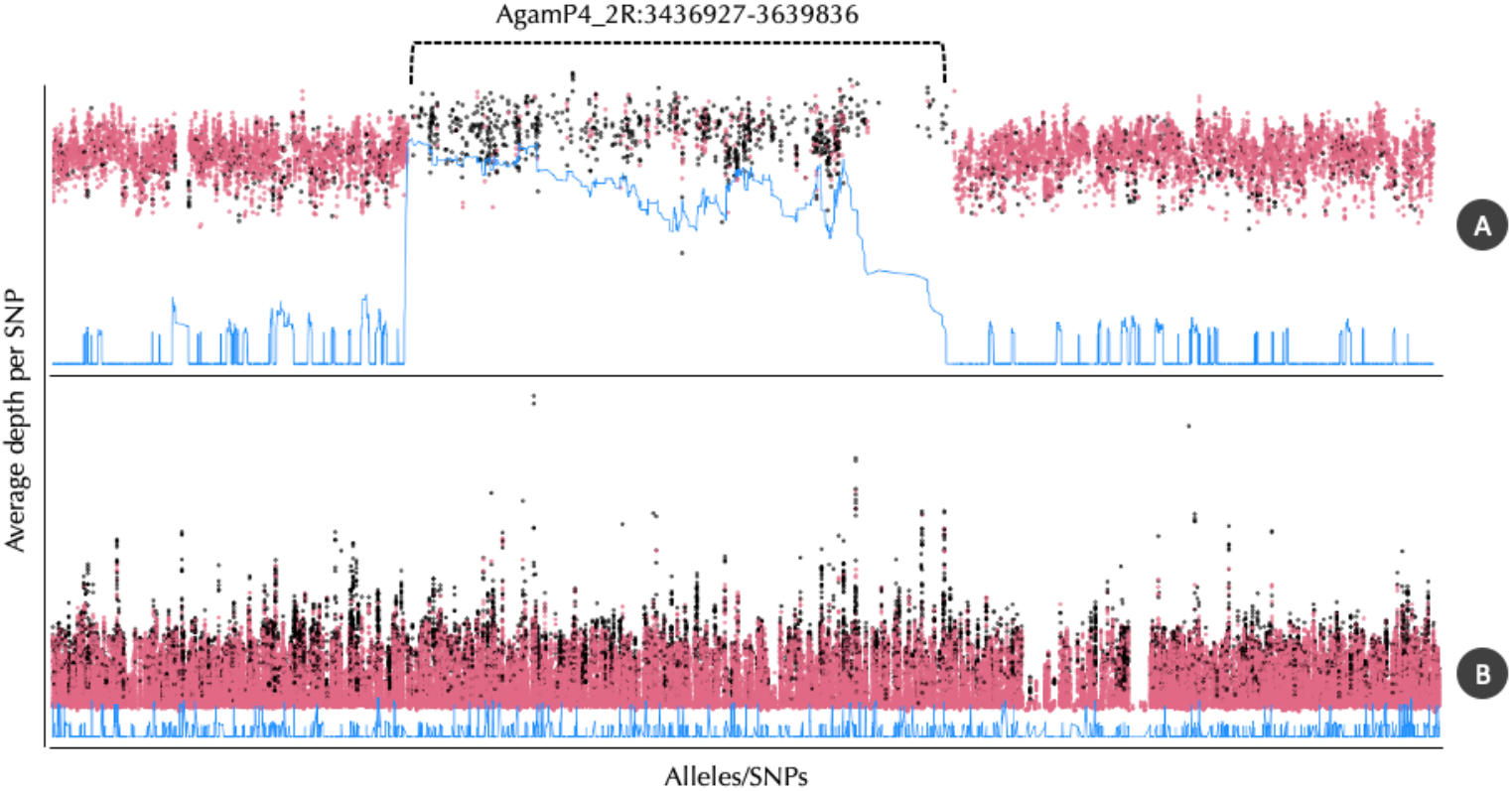
The average read coverage depth variation of alleles sequentially arranged on the genomes of A) *Anopheles gambiae* and B) Norway Spruce. Black points indicate the detected putative CNVs by rCNV; the blue line shows the moving window average of the depth values. AgamP4_2R:3436927-3639836 is the previously recognized region of duplication in the right chromosome arm of *Anopheles gambiae*.

For all datasets, we observed higher density distribution of average read depth coverage in SNPs located in putatively multi-copy regions (Fig. 4). Since we did not use average read depth coverage in any metrics, this further supports our classification. Nevertheless, different levels of overlap can be seen in the density distribution in Figure 4. In addition to factors that were not considered in the analyses (e.g. sequencing errors, local variation in coverage), two main factors can explain the overlap: 1) SNPs misclassified as putatively located in multi-copy regions or the opposite, and 2) among SNPs putatively located in multi-copy regions some are located in CNVs which are, by definition, polymorphic for the number of copies in the population. The latter is effectively illustrated in the *Anopheles* mosquito dataset. Furthermore, the local enrichment of putative CNVs in specific genomic regions where the average depth coverage is higher (i.e., peaks) for the Norway spruce and mosquito datasets (Fig. 3B) further supports a low false positive detection rate and corroborates the variation in the density distribution of the average dead depth coverage in putative CNVs.

**Figure 4.**
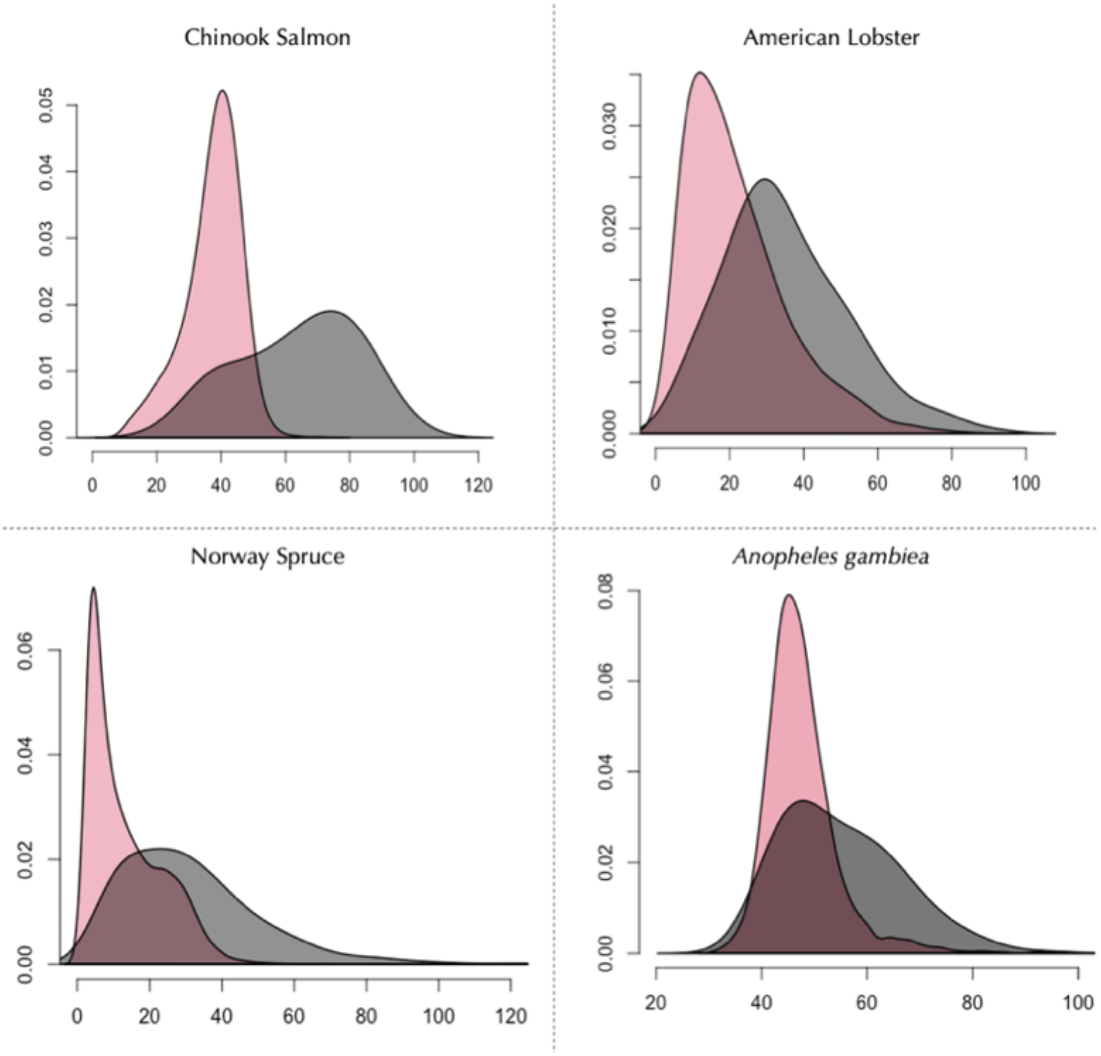
The density distribution of average read depth of putative CNVs and non-CNVs in the studied SNP datasets (red: non-CNVs, black: putative CNVs).

## Discussion

Efficient and more accurate detection of SNPs located in the multi-copy region from genomic data can immensely improve genetic analyses by i) generating a clean dataset free of SNPs that are in multi-copy regions and low confident SNP calls due to technical issues, and ii) detecting putative CNVs and their relative copy numbers that can be used in quantitative and population genomic studies. Generating such datasets with minimal errors is almost impossible without whole genome sequencing data and well-annotated reference genomes. Yet, the vast majority of genetic data consists of SNPs datasets generated through genotyping by sequencing or target sequencing methods. With our framework, we provide a robust filtering method for SNPs showing unexpected patterns in allelic ratios (e.g., because of sequencing bias, CNVs/paralogues, or sequencing errors) and a robust CNVs candidate detection pipeline, both from single nucleotide polymorphism data. Our framework is user-friendly, easily accessible, and low demanding of programing skills as it is implemented within the R infrastructure. The approach implemented in the *rCNV* package is statistically sound, and the results tested in our case studies show substantial accuracy, especially with high-throughput non-whole genome sequencing data.

### Thresholds and false positive detection propensity

One of the biggest challenges in CNVs detection using SNPs data is the propensity for false detection due to, among other factors, sequencing, and probe-biases (see examples in chapter 4 of Neves, 2013), as well as sampling effect due to large variations in read depth and the number of heterozygous samples, which can introduce deviation from expected patterns (Figs. 1 and S1). For a given SNP, what should be considered a significant deviation from the expected allelic ratio, would depend on both the number of heterozygotes and the depth of coverage across the heterozygotes. We thus recommend eliminating SNPs with a low number of heterozygotes (< 4) and low average read depth (< 5) (the default parameters in *rCNV*) in CNVs detection. With the function *depthVsSample* in the *rCNV* package, it is possible to visualize how the confidence intervals around a given expected allele ratio vary with varying numbers of heterozygotes and depth of coverage values (Fig. S1A) and the function *power*.*bias* demonstrates the 95% confidence level *Z*_*X*_ based detection power of allele biases for a given number of samples for a given range of depths (Fig. S1D-E). In our R package, however, the users are given the choice to adjust these parameters to best fit their data. Moreover, outlier samples (i.e., samples with extremely high or low total depth values) resulting from sequencing anomalies can contribute to the classification of SNPs by skewing *Z*_*x*_ and 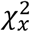 calculations. Outlier samples also have a significant effect on the normalization factor that we calculate to normalize read depth. We observed this in the American lobster dataset; false positive detection was higher when the outlier samples were not removed.

In the *HDplot* method developed by McKinney et al. (2017) and further implemented by Dorant et al. (2020), single threshold values are defined through visual observation of the distribution of various statistics (e.g., the median allelic ratio in heterozygotes and excess of heterozygotes in both, the proportion of rare homozygotes or inbreeding coefficient only in Dorant et al. 2020). For a given statistic, the same threshold is thus applied to all SNPs. This approach can serve as a preliminary assessment or as a standalone analysis if prior knowledge about ploidy level and/or paralog status is present. However, it overlooks the continuity in deviation from the expected allelic ratio due to variations in the number of heterozygotes and depth of coverage from one SNP to another, polymorphism in the number of copies, and/or variation in frequency in the dataset as illustrated in Fig. 1 and S1.

In our framework, we have addressed these shortcomings by computing two statistics, *Z*_*x*_ and a 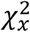, allowing us to test the significance of the deviation from the expected allele ratio for each SNP independently. It allows taking into account SNPs-specific parameters (e.g., number of heterozygotes, depth of coverage, capture bias) thereby minimizing false positive detection. Still, the users have the flexibility of adjusting the thresholds for the *p*.*value* to better suit their data as well as the stringency for the false positive detection rate. Note also that levels of stringency for the “*deviant*” SNPs detection may affect the detection of the SNPs putatively located in multi-copy regions using the *intersection-set* approach. Therefore, we also provide the user with a threshold-free approach using *K*-means clustering. We obtain highly overlapping classifications using both methods, specifically for RAD-seq data, where reference bias (in a broad sense) is minimum compared to probe-based methods (e.g., exome-capture).

### Deviants and putative multi-copy regions

It is a fair evaluation that the methods used in our framework may not result in the same accuracy for all data sets. In fact, the accuracy is heavily influenced by the quality of the data as the reliability of SNP calling and read depth analysis are influenced by the quality of the raw sequences (Barbitoff et al., 2020; Borges et al., 2020; Gabrielaite et al., 2021). With real-world data, we show in our detection of *deviants* and multi-copy regions, that our framework is efficient at encapsulating both undesired loci (deviants) to be removed in SNPs-based genetic analyses and important structural variants (CNVs or paralogues) for downstream analyses. Further, depending on the quality of the data (e.g., probe bias) and the purpose of downstream analysis (e.g., with CNVs as a focus, or excluding multi-copy regions), stricter filtering criteria in the deviant detection will eliminate false positives from the putative duplicates. Given our simulations and power analysis, we are confident in the fact that the rCNV approach is reliable enough to separate false positives/negatives from deviants provided that the underlying data is robust enough (in terms of depth and number of heterozygotes). We provide the user with the *power*.*bias* function in the R package to test this on simulated data. We have high confidence that the appropriate use of filtering criteria (i.e., *test* and *filter* argument of the *cnv* function in the *rCNV* package) will minimize the false positive detection of multi-copy regions from deviants. Finally, if one wants to discriminate fixed paralogous from actual CNVs, we recommend using a combination of the proportion of heterozygotes, which should be close to one, and the coefficient of variation that should be low as no difference of coverage would be expected from one individual to another, regardless of the so-called “apparent heterozygosity”.

### Different sequencing technologies

While DNA sequencing technologies are rapidly improving with better quality and more accurate generation of sequences, most non-whole genome sequencing techniques carry certain degrees of error and anomalies (Jones & Good, 2016; Krumm et al., 2012; Meynert et al., 2014). Studies show that exome-capture technologies are approximately 95% accurate at detecting the alternative allele at a mean depth of 40 reads compare to WGS because of probe sequencing bias (Lelieveld et al., 2015; Meynert et al., 2014). Generally, we refer to as *probe-bias* when the (exome-capture)-PCR probes have a higher tendency to capture reads from one allele of the same locus than the other alleles. Among many factors, the GC content, DNA concentration, and annealing temperature are the most commonly known causes of such biases (Barbitoff et al., 2020; Borges et al., 2020; Neves, 2013). While these differences can be dramatic, most importantly, the captured reads for two regions and two alleles in a heterozygote can be significantly different with probe bias, leading to a significant deviation from the theoretical allele ratio of 0.5, and therefore, can lead to false detection of CNVs. We show this tendency with the Norway spruce exome capture dataset. Although we have addressed this issue in our framework by assessing the observed allelic-ratio across all samples at a given SNP (i.e., also taking into account the ratio between homozygotes), we recommend that, if a dataset is more prone to probe bias and there is prior knowledge of the bias ratio, the user input that information when detecting deviants and multi-copy regions (e.g., *test=z*.*all/chi*.*all* option in *dupGet* and *cnv* functions). Nevertheless, one should keep in mind that in cases where poor sequencing and high probe bias are prevalent, the detection of multi-copy regions may be less reliable as seen in the simulations (see Fig. S1) when the number of heterozygotes and the median allele depth are low.

### Prospects

#### Further annotation of CNVs

When the SNPs are ordered along a reference, we provide the user with the possibility to use a density-based sliding window approach to further refine the list of SNPs putatively located in CNV regions. Ideally, if a locus is truly located in a multi-copy region, close-by SNPs should also be classified as *deviants* while deviation caused by sequencing error should be more scattered along the genome. The function *dup*.*validate* in the rCNV package is dedicated to detecting regions enriched for deviant SNPs within a sliding window along a chromosome, scaffold, or a sequence of any given length. Users can define both, the length of the window and the threshold above which a locus is considered a *true deviant*. We demonstrate this with Norway spruce and *Anopheles* mosquito datasets where deviant SNPs with a frequency threshold less than 0.2 within a moving window of 10,000 are flagged as low confidence. This step of filtering is based on the assumption that at least 20% of the loci within a duplicated region must be detected as deviants/CNVs for the loci in the respective region to be considered a deviant/CNV of high confidence.

#### Relative Copy Number (RCN) and Environmental Association Analysis (CNV-EAA)

The depth values of putative CNVs can be used for population genetic analyses such as *V*_*ST*_ or multivariate and ordination analyses such as PCA to examine the population structure and local adaptation using CNV (e.g., Fig. S8 and S9). We show this application with the American lobster data; to compare our output with Dorant et al. (2020) results, we performed two downstream analyses; the ordination of putative CNVs from the lobster data (Fig. S8) showed a strong spatial structure among individuals that is not explained by the geographical distribution but rather by the distribution of Sea Surface Temperature (SST) as was observed by (Dorant et al., 2020a). Further, variance fixation index (*V*_*ST*_) analysis using the normalized depth values of detected putative CNVs showed a correlation to *F*_*ST*_ calculated with the genotypes of the non-CNVs (putatively non-duplicated) in the analyzed data set (*r*^*2*^= 0.82), yet the population structure obtained from CNVs was much stronger than that obtained from non-CNVs. However, categorizing the samples or populations into relative copy number variants (RCN) will allow capturing a different set of genetic variations that is more meaningful for population structure and adaptation analyses. Further, RCN can be incorporated into environmental association analysis (EAA) since the relative copy number can capture quantifiable genetic variation to adaptation (North et al., 2020; Prunier et al., 2019). However, correctly deciphering the RCN from the depth values is challenging, especially when the depth values vary continuously among samples. Clustering methods (e.g., Fraley & Raftery, 2002), breakpoint analysis (e.g., Chong et al., 2017; Faust & Hall, 2012), and Bayesian criterion (e.g., Fraley & Raftery, 2007) are potential candidates for categorizing such data. These methods however do not determine the relative copy number in accordance with the depth increment. For instance, clustering methods may assign an RCN value of one (RCN=1) for a depth value of 50 and two (RCN=2) for a value of 150. However, the correct relative copy number for such an increment should be three (RCN=3) as the second sample has three times the read depth value as the first one. Therefore, the development of a statistical method to correctly categorize the relative copy number from the depth values is underway for rCNV.

One major use of CNV from large-scale population genetic analyses is to detect spatial genetic variation and local adaptation. Environmental association analysis (EAA) with currently available methods (see Manel & Holderegger, 2013) relies on the allele frequency calculated from genotypes to assess the genetic variation corresponding to the environmental variables. However, allele ratios calculated with depth values in duplicated loci are not representative of the actual copy number variation among individuals as two different copy numbers can produce the same allele frequency. An alternative method used in previous studies (e.g., Cayuela et al., 2021; Dorant et al., 2020) is to use the composite values of a PCA performed on the normalized total depth values of the duplicates in a multivariate analysis with the environmental variables. Nevertheless, the use of normalized depth values undermines the relative variation among populations and individuals. Therefore, the relative copy number (RCN) will provide a comparative analysis with local adaptation.

### Benchmarking rCNV

To compare the efficiency of detecting CNVs from SNPs data using the rCNV package, we explored the scientific literature and the major bioinformatics platforms to find comparable methods and software. Despite the availability of a range of methods and software for detecting CNVs from whole genome sequences (see Gabrielaite et al., 2021), we found only two methods comparable (see below) to some aspects of our approach. While there is an array of methods developed for detecting CNVs from short-read sequencing such as read-pair split-read (SR), read-pair (RP), and assembly (AS), algorithms based on read coverage differences in the genome, also referred to as read-depth (RD) is the most common method of CNV detection in most software packages available (Gabrielaite et al., 2021; Kosugi et al., 2019). However, most if not all, such algorithms rely on complete and precise reference genomes (e.g., GATK gCNV, ExomeDepth) or require whole exome capture or whole genome sequences (e.g., CNVnator, LUMPY). Further, the raw data used in each software package is specific to their own method (e.g., Stacks software -Catchen, Hohenlohe, Bassham, Amores, & Cresko, 2013; Rochette, Rivera-Colón, & Catchen, 2019) or requires a great deal of computation power and memory (e.g., CLC Genomics Benchmark – fastq; cnMOPS – bam files) (see Gabrielaite et al., 2021 for a detailed comparison). In contrast, while most available methods are dedicated to exhaustive exploration of structural variants including CNVs from WGS data, our framework is ideal when such methods are not available, for instance for studies involving non-model organisms. Finally, most of the available software/methodologies require at least an intermediate level of programing skills. Therefore, in essence, our framework and *rCNV* package stand out from these methods in three main aspects: 1) use of VCF files with SNP calls, which are relatively small in size compared to other input data formats, 2) not requiring whole genome sequences or precise reference genomes, and 3) the detection is statistically robust and does not demand expert programming skills. Overall, the framework implemented in rCNV is reliable, cost-effective, easy to use, and significantly fast.

In addition to the robust and easy detection of CNVs, *rCNV* offers effective pre-processing of the data (e.g., filtering of SNPs based on missingness, heterozygosity, relatedness, and outliers) as well as basic post-detection downstream analyses such as V_ST_ calculation, RCN, and sliding window-based CNV validation.

#### Comparison with the HDplot method (McKinney et al. 2017)

The method proposed by McKinney et al. (2017) aims at identifying SNPs where median allelic-ratio (ratio between the depth of coverage of two alleles at a given position) in heterozygotes individuals deviating from 1:1 (expected ratio in heterozygote when both alleles are as much covered) as well as the SNPs with excess of heterozygotes. This method was developed to filter out SNPs located in paralogous regions and it can have limitations when used to call CNVs. First, it is a threshold-based approach reliant on the expectation that allelic ratio can be drawn from a pre-determined number of copies. For CNVs however, the allelic ratio variation is more continuous as it depends on the number of copies in a given individual and their frequencies across all individuals; two values that can vary from one locus to another. Second, despite being effective as a priori, the use of median and a threshold is sensitive to sampling effect (variation of depth and number of heterozygotes among SNPs) that can lead to a high false-positive detection rate of CNVs, especially when removing putative paralogous. Finally, sequencing technologies are prone to errors, especially, widely used target capture technologies such as exome-capture and SPET where capture biases can occur toward the reference allele (see Lelieveld et al., 2015). In such cases, the expected ratio between the reference and alternative alleles is no longer 1:1 and should instead reflect the capture bias. *rCNV* has addressed such shortcomings by statistically scrutinizing the probe bias and implementing methods to correct for them while accounting for the sampling effect inherent to depth variation and the number of heterozygotes between SNPs.

#### Comparison with the stacks_workflow method in Dorant et al. (2020)

We compared the stacks_workflow (https://github.com/enormandeau/stacks_workflow) used by Dorant et al. (2020) for a comparison of a similar detection method to *rCNV*. The method implemented in stacks_workflow is, in essence, the same *HD-*plot approach developed by McKinney et al. 2017 and visual observation of the distribution of additional statistics (e.g., F_IS_, the proportion of rare allele homozygotes) with additional filtering steps to eliminate sampling bias. Despite the additional filtering and more statistical-based classification of paralogs, this workflow still suffers similar drawbacks as mentioned above. To test the efficiency of the pipeline, we parallelly followed their workflow steps with our methodology (as in Fig. 1) on the American Lobster data set. From filtering raw VCF files to the detection of putative duplicates, Dorant et al. (2020) used three Python scripts and two R scripts, which altogether accounted for a total of 12 minutes and 23 seconds for the complete analysis. However, this time is without the time spent on visual observation of the various allelic ratio-based statistic plots they used to determine the thresholds for sorting duplicates from “singletons”. In contrast, the *rCNV* package only took 31.01 seconds to complete the detection process with a statistic-based setting of thresholds. Therefore, compared to the method used in the stacks_workflow, *rCNV* is not only computationally faster but also detects duplicates based on a robust statistical approach with minimal intervention from the user. Further, several Python scripts used in stacks_workflow are specific to VCF files generated from the Stacks software package for calling SNPs and are usable with only RADseq data. Moreover, in the rCNV workflow, we employ various pre-processing steps, including additional normalization methods, odd read depth correction and reassignment of false genotype classification.

One main purpose of the rCNV framework is to minimize the bias introduced by CNVs/paralogs in population genetic estimates when poorly accounted for as they tend to introduce “apparent heterozygotes” (i.e., pseudo-SNPs). In that regard, being blind to CNVs/paralogs that have not yet diverged where all the copies are strictly identical, should not cause false detection as they do not introduce bias in apparent heterozygote frequencies. Note that the same rationale holds for CNVs segregating at a low frequency just enough to be not detected; for instance, if the number of heterozygotes is low. However, it is more problematic in the latter case with the rCNV framework, which is to detect putative CNVs and use them as “genetic markers”. In that specific case, rCNV would indeed fail to detect these CNVs/paralogs. Finally, to avoid including false positives in downstream analyses, i.e., SNPs erroneously classified as putatively located in a multicopy region, we strongly encourage the users to make use of the various statistics we generate. CNVs, by definition, are expected to vary in copy number between individuals, which should be reflected in the coefficient of variation we output in allele information. Also, CNVs often span over several SNPs and one can expect a consistent signal for multiple SNPs located close by. Density-based approaches such as the “window-based approach” suggested in our framework can be used to further filter the “CNVs” dataset. Furthermore, our simulations and power analyses (Fig. S1A, D&E) clearly demonstrate the limitations to unequivocally classify SNPs into any category when the depth and/or number of heterozygotes are low without prior knowledge of the copy number status. Also, given the shape of the site frequency spectrum, the fact that CNVs are expected to be less frequent than “regular” SNPs and that CNVs/paralogous sequences tend to induce so-called “apparent heterozygotes”, one can expect most of the SNPs with a low number of apparent heterozygotes to be located in single-copy regions.

## Acknowledgments

We thank Swedish National Infrastructure for Computing (SNIC) for resource allocation for high-power computing under the project numbers SNIC 2021/6-313 and SNIC 2021/5-540. The Nilsson-Ehle grant by the Royal Physiographic Society of Lund awarded to Piyal Karunarathne (Grant No. 42191) was immensely helpful for acquiring the resources to test the features of the R package locally. We wish to thank Freja Lindstedt for her generous work in further testing the CNVs detection on real-world data.

## Data Accessibility and Benefit-sharing

### Data Accessibility

All the genetic data used in the paper are mentioned in the text with their references where applicable. All the scripts and simulation outputs can be accessed from the GitHub repository (<u>link</u>)

### Benefit-sharing

Not applicable.

## Author Contributions

The framework design was conceived by PM; the methods and statistical approaches developed by PM, PK, and QZ; PK developed the software package and analyzed the data. KS improved and tested the functions and PM and QZ beta-tested the R software package. PK wrote the manuscript with input from all the authors

