## supplementary data for "A new framework for detecting copy number variants from single nucleotide polymorphism data: ‘rCNV’, a versatile R package for paralogs and CNVs detection"

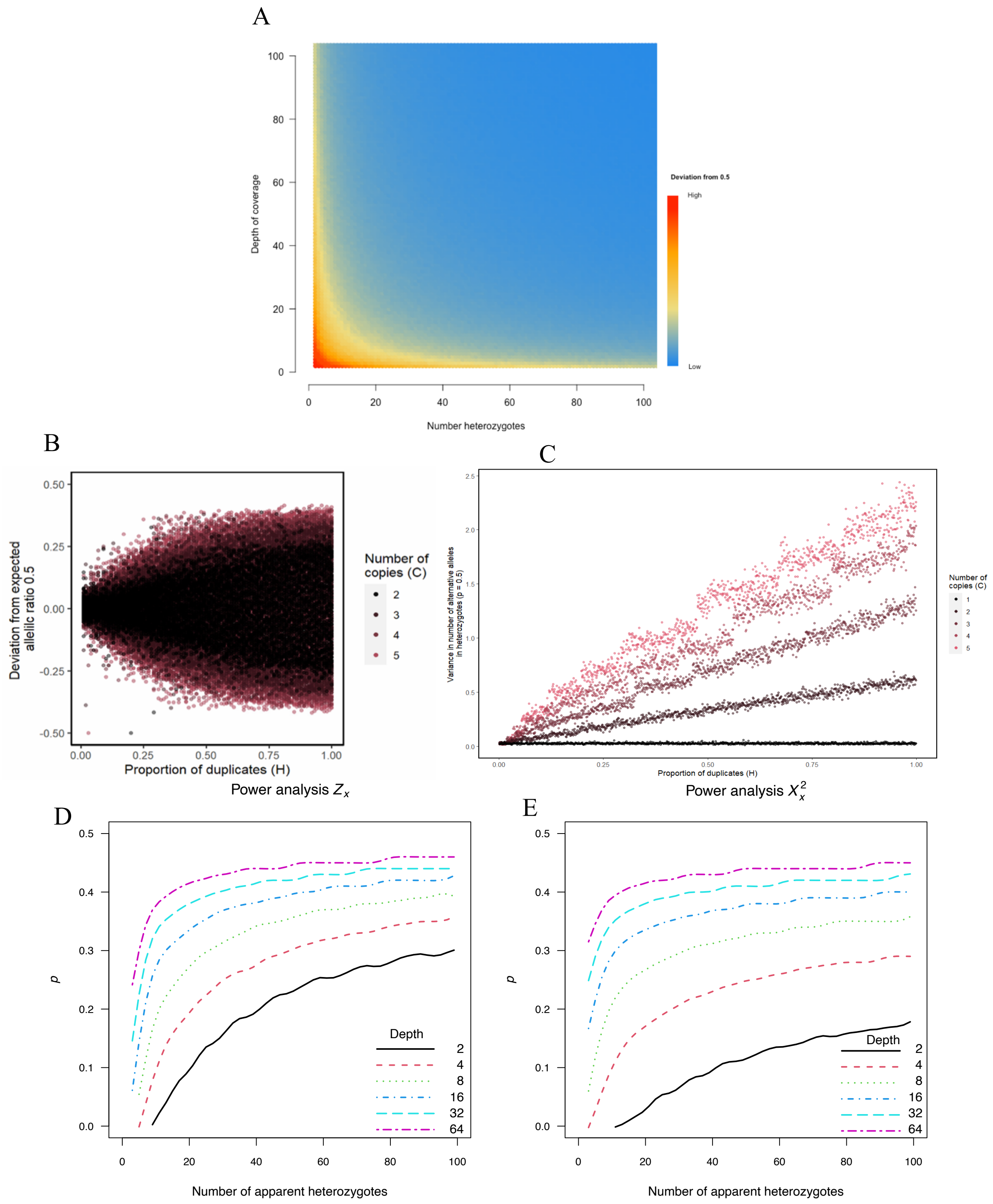


**Figure S1. Patterns of allelic ratio variation and power analyses.** Deviation from expected allelic ratio because of **A.** sampling effect introduced by both sequencing depth and number of heterozygous genotypes for a given SNP and **B**. Number of copies and proportion of segregating duplicated allele. **C.** gives the variance in number of alternative alleles as a function of number of copies and proportion of segregating duplicated allele. Note that for NC = 1, the x axis is the proportion of heterozygotes. **D** and **E** are power analyses of detection of significant deviation in mean ($Z_x$) and of higher variance ($X_x^2$) than expected as a function of number of heterozygote individuals ($n$) and read-depth (N). For details, please refer to main text Material and Methods §"Simulations".

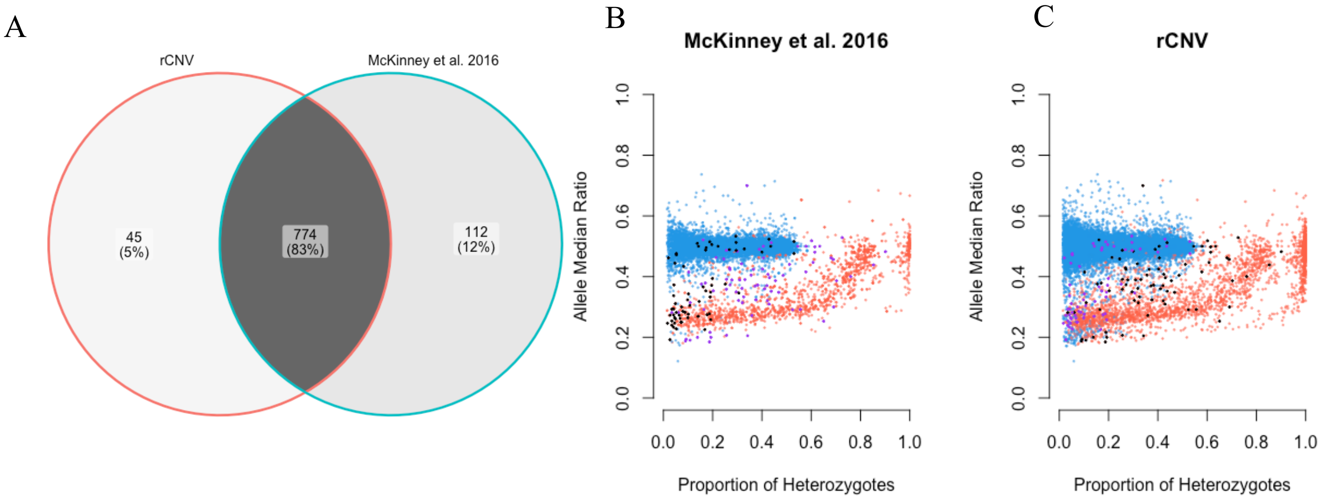


**Figure S2. Comparison of CNVs detection in Chinook salmon by rCNV and McKinney (2016):** A – Overlap of putatively duplicated alleles for the two methods, B, C – overlap of pseudo false positive and pseudo false negatives (red – duplicate/deviants, blue – singletons/non-deviants, black – *pFP*, purple – *pFN*)

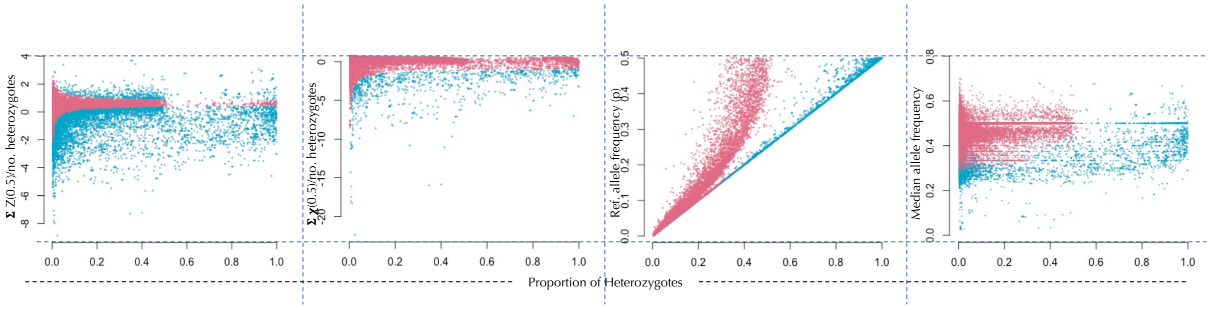


**Figure S3. Different ranges of deviant SNP detection:** by (plots left to right) Z-score distribution of depth values, chi-square distribution of depth values, excess of heterozygotes, and the K-means clustering on American lobster data (Dorant et al. 2020)

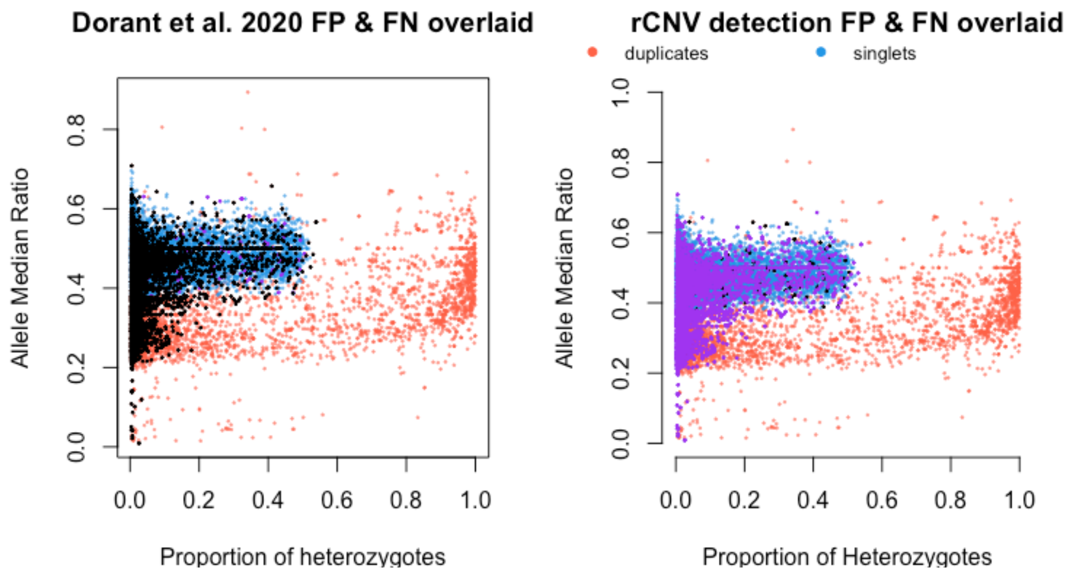


**Figure S4. Overlap of false positive and false negative detections by rCNV and Dorant et al. 2020.** Black – pseudo false positive, purple – pseudo false negative

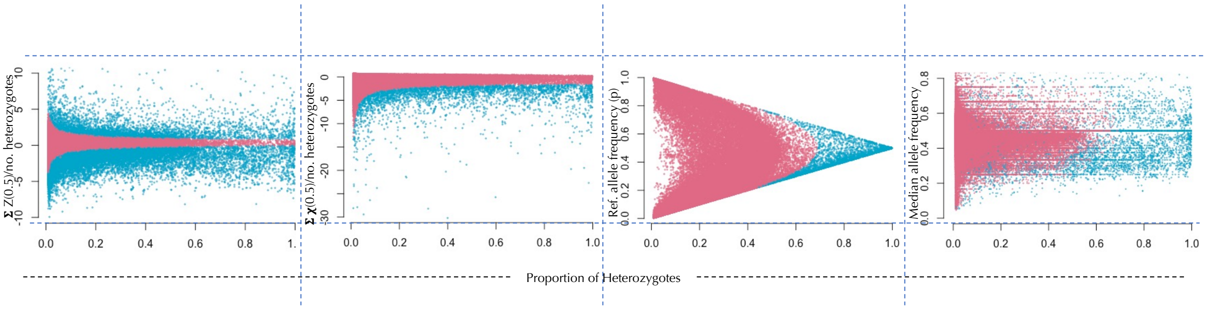


**Figure S5. Different ranges of deviant SNP detection:** by (plots left to right) Z-score distribution of depth values, chi-square distribution of depth values, excess of heterozygotes, and the K-means clustering on Norway spruce data (subset from Chen et al. 2019).

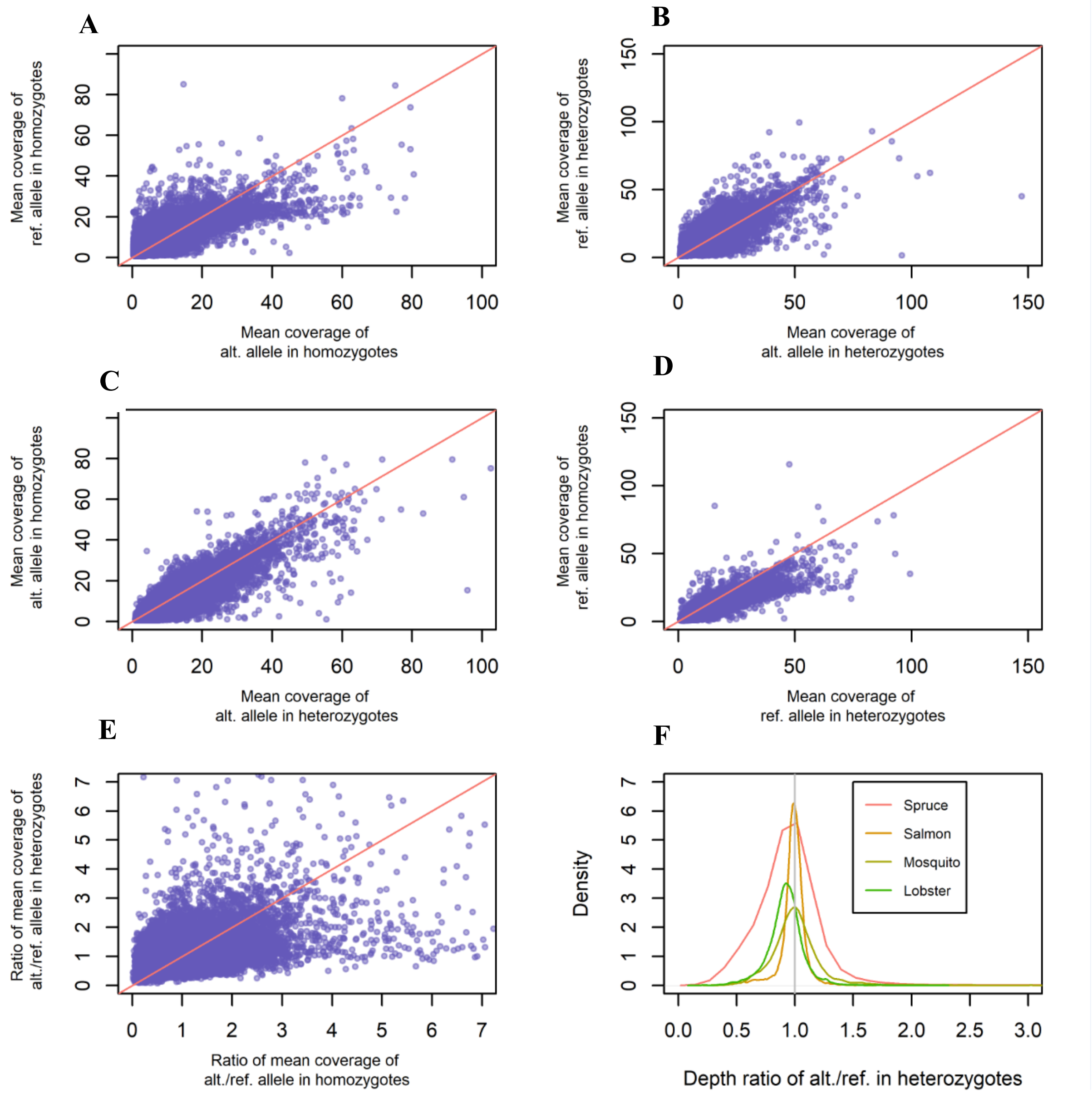


**Figure S6: Investigation of probe/reference bias: Plots A to E are** correlations between coverage values of reference and alternative alleles in Norway spruce between various genotypes in Norway spruce (Red lines are identity line y=x; $r^2$: 0.78, 0.80, 0.87, 0.87 and 0.21 A-E respectively). Plot F shows the distribution of allelic ratio for the four datasets.

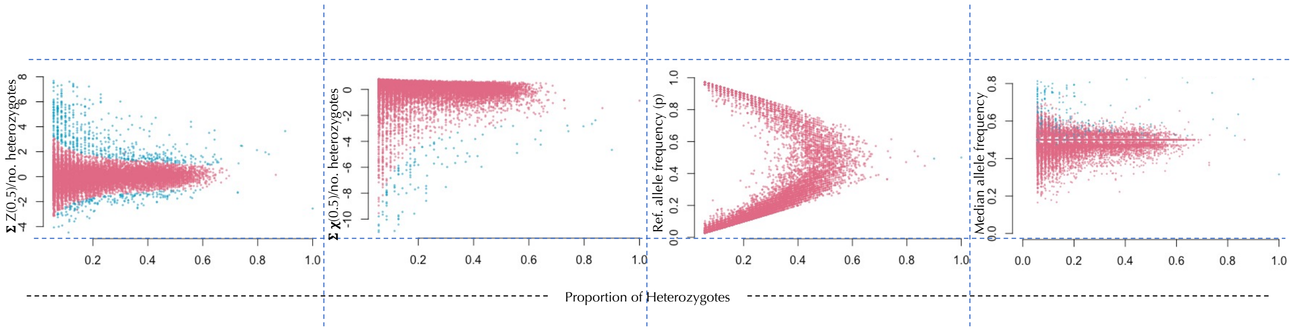


Figure S7. **Different ranges of deviant SNP detection in *Anopheles gambiae* data**: by (plots left to right) Z-score distribution of depth values, chi-square distribution of depth values, excess of heterozygotes, and the K-means clustering (pink – non-deviants, blue – deviants).

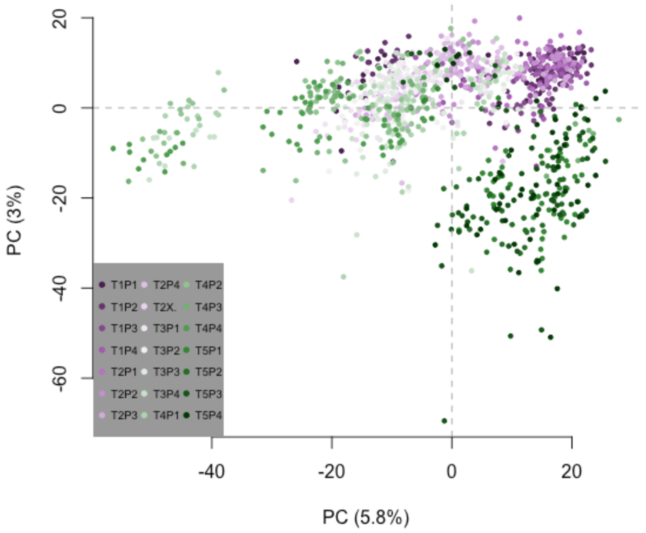


**Figure S8. PCA of normalized allele depth values of putatively CNV sites of the American Lobster dataset (data - Dorant et al. 2020).** The color scale shows different populations collected from different locations.

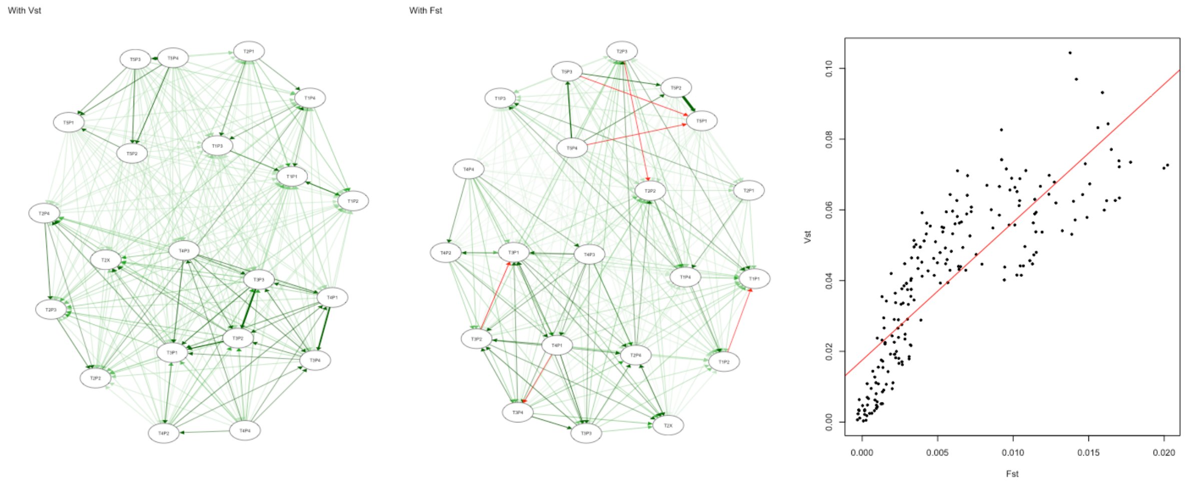


**Figure 9. Comparison of FST allele frequencies and VST computed from depth values for non-CNVs and CNVs respectively. A, B**. Similarity/dissimilarity igraph network plots; **C**. Regression of *Vst* vs *Fst*.

**Table S1. Recommendations for using different options in important functions included in rCNV**

| Step/function | Description | Recommendation |
|---|---|---|
| Normalization | Trimmed means of M-values (*TMM*) was proposed by (Robinson & Oshlack, 2010) calculates the scaling factors for individual samples, using which the effective library size can be calculated and thus normalized depth values. We adopted median ratio normalization (*MedR*)and quantile normalization (*QN*) methods from normalization methods for differential gene expression analysis of RNAseq data (Maza, Frasse, Senin, Bouzayen, & Zouine, 2013). The PCA normalization method assumes that the variation of samples from a principal component analysis is due to library size and removes the top PCs following a modified Kaiser's rule (eigen value < 0.7). | Although all normalization methods used in rCNV does a fairly good job in adjusting the batch effect/sequencing anomalies, the method of choice depends on the spread of the library size. Therefore, it is recommended to review the sample library size before normalization. If the user cannot decide on a method, median ratio (MedR) is a safe method. |
| *dupGet*/deviant detection | *dupGet* function efficiently combines deviant detection and allows users to use either Fisher's exact test or Chi-square test to determine the SNPs with excess of heterozygotes. With deviation from expected allelic ratios, users are given the options to use the Z-score values with or without probe biases (i.e., z.all or z.05 respectively) and the chi-square test on either heterozygotes (*chi.het*) or all (including homozygotes) (*chi.all*). | The choice of statistics for deviant detection should be based on probe bias. Putative probe biases can be assessed using the plots generated in the *allele.info* function. |
| Putative CNVs/ intersection set | If the underlying data was generated with sequencing methods that are less prone to probe-bias, such as RADseq, the excess of heterozygotes in combination with either *Z*-score or $X^2$-score for allele occurrence probability $p$ = 0.5 will filter the optimal number of putative duplicates from the deviant SNPs. On the other hand, the sequencing technologies such as exome capture are more prone to sequencing bias and thus Z-score and chi-square tests with $p$ determined on the distribution of allele frequency should be used in the filtering steps. | We recommend using both a less stringent approach such as $p$ = 0.5 and more stringent z.all and chi.all for smaller datasets to compare the different. |
| Putative CNVs/ K-means clustering | The K-means clustering is independent of threshold or cutoff values for clustering the duplicated and non-duplicated SNPs and hence unsupervised. Since Z-score, Chi-square and delta are calculated on the read-depth values of alleles in heterozygotes, clustering can filter CNVs based on read-depth variation without being sensitive to extreme read-depth values. In contrast, Z-score and Chi-square, despite being highly effective in detecting deviants especially at lower depth values and lower heterozygosity, can produce false positives and false negatives due to strict threshold values. However, K-means clustering lacks power at higher heterozygosity in combination with lower read-depth values. Therefore, we combine excess of heterozygosity and K-means for filtering putative CNVs. | Although K-means clustering is always a safer method for filtering, we recommend the user pay attention to the proportion of heterozygosity and the distribution of depth values to avoid false negatives. Intersection set performs better for datasets with lower depth and higher proportion of heterozygotes (e.g., mean depth < 10 & propHet > 0.7) |

*Comparison of rCNV to other R packages*

We found only one more R software package that implements a similar approach to detecting CNVs as rCNV. The R package “radiator” (Gosselin, 2020) is dedicated to the exploration, manipulation, visualization, imputation, and exportation of RADseq and GBS data. The current version (v.1.2.2) is only hosted on GitHub (https://github.com/thierrygosselin/radiator/) and indicated to be “maturing.” The function *detect.paralog* in “radiator” has been developed to detect duplicated loci from SNP data using the *HDplot* method described by McKinney et al. (2017). However, we were not able to test the “radiator” package or the *detect.paralog* as the examples provided in the function itself do not work and the author of the package has not provided any other alternative to test the function as it only accepts data formats internal to the “radiator” package. Our attempts to contact the author were not successful either. Overall, the “radiator” package is not usable for detecting CNVs in the current state.
